# Quantitative modeling reveals sources of variability in transcriptional activation assays

**DOI:** 10.64898/2026.01.30.702786

**Authors:** Mark Greenwood, Kenneth F. Reardon, Ashok Prasad

## Abstract

Reporter cell assays, such as those used to detect estrogenic chemicals, can detect target chemicals at low concentrations and can be used to analyze chemical mixtures without *a priori* knowledge of the mixture components. However, the outputs of these assays are affected by biological variability, which complicates their interpretation. Here, we describe and demonstrate a workflow that is useful for determining potential sources of biological variability and optimizing the performance of cell-based assays. The workflow involves developing an appropriate mathematical model for a transcriptional activation assay, calibrating it with experimental data, and conducting sensitivity analysis to characterize individual components of the genetic circuit based on their effect on the reporter signal output. This workflow was tested using an estrogen receptor transcriptional activation assay. For this circuit, our analysis predicts that controlling estrogen response element number, promoter strength, and reporter signal degradation rates minimizes reporter output variability. We show that careful model development, calibration, and analysis can offer biologically relevant insights to minimize the variability of cell-based assays and improve genetic circuits for increased sensitivity and dynamic range.

## 1. Introduction

Estrogen disrupting chemicals (EDCs), including estrogenic chemicals, have significant negative consequences for human health and the environment (1,2). Since the majority of the thousands of chemicals in use have not been rigorously tested for EDC activity, and new chemicals are being introduced every day, accurate and rigorous tests for EDC activity are needed (3). Cell-based assays are often used for screening EDCs. These are typically transcriptional activation assays, designed to test for xenoestrogens by linking an optical luciferase-based readout to estrogen receptor activation in human cells (4–8). These assays rely on the binding of the suspected xenoestrogen with the estrogen receptor and measure its capacity to activate genomic estrogen signaling of a reporter gene expressing luciferase. Cell-based assays that interrogate EDC activity do not yet replace animal models due to the complex interactions of EDCs with endogenous hormone response (9). Nevertheless, cell-based assays are ideal for identifying potential hazards from large chemical libraries, making them indispensable for applications such as supporting regulatory efforts with dose-response data and screening chemical mixtures (3).

Cell-based assays for detecting EDCs are part of a broader group of reporter assays that are widely used for drug testing and toxicology (10–12). These assays can detect target chemicals at low concentrations and evaluate complex mixtures without prior knowledge of their individual components or interactions. However, their readouts are inherently influenced by biological variability, which can limit precision and reproducibility (13). A general phenomenon, cell heterogeneity is a major factor affecting the reproducibility of experimental results (14). In one study, fluorescence levels measured by different laboratories using the same genetically engineered constructs in *Escherichia coli* ranged over an order of magnitude (15,16). Cancer cell lines display more variability due to a high mutation rate and genomic instability (17). Protein expression levels vary from cell to cell (18), leading to differences in parameters and reaction rates on a cell-to-cell and experiment-to-experiment basis. By reducing cellular variability, cell-based assays can be improved.

Advancements in synthetic and systems biology have transformed the potential capabilities of cell-based assays, developing techniques to specifically design circuit architecture to tune assay attributes such as sensitivity, dynamic range, and robustness (19). These techniques have led to improved sensing circuits and the ability to resolve underlying molecular mechanisms (20,21). One of the key ingredients in modern synthetic biology approaches is its reliance on forward engineering, i.e., mathematical modeling, quantitative testing, and characterization of the sensing circuit, and its use for optimization of circuit design and performance (22–25). In addition, global sensitivity analysis has been demonstrated as a computational tool to predict the effects of mutations on a genetic circuit in *E. coli* (26). While forward engineering has been demonstrated in mammalian cells, these methods have yet to be applied to reporter gene circuits used in cell-based assays.

Here, we demonstrate how forward engineering methods from systems and synthetic biology can be applied to reporter gene circuits to optimize performance and identify sources of variability. Using a reporter gene circuit for EDC cell-based assays as an example, we first develop a mathematical model and show that robust parameter estimation can be performed even with limited data. We then analyze the mathematical model using global sensitivity analysis to identify the parameters that most affect the reporter output. We propose that parameters that lead to the most significant quantitative changes in dynamic range and sensitivity, both of which are important performance attributes for reporter assays, are the parameters that would also contribute the most to variability. Strategies to control these parameters would minimize cellular variability and optimize circuit performance with respect to user-defined specifications. New circuit designs, based on advances in synthetic biology, can then be implemented.

## 2. Materials and Methods

### 2.1. Model development and assumptions

The sensing circuit model was developed assuming that the law of mass action governs reaction kinetics between constituents of the model. The reactions and model equations are presented in the supplementary material (Equations S1 and Figure S1). Estrogen receptor (ER) activation is complex and dependent upon both ligand and cell type, relying on ERα66, ERα46, ERα36, ERβ isoforms, GPER/GPR30, and membrane-bound ER along with EGFR (27–29). Thus, the model was simplified with several assumptions. Ligand is assumed to be instantaneously diffused in the cell. Only ERα66 is assumed to drive transcriptional changes in the reporter gene. Basal expression is assumed to be attributable to ER binding in the absence of ligand. ER dimerization is lumped. Ligand-based activation is modeled by a change in the binding rates of ligand-bound ER. Transcription and translation are governed by first-order kinetics.

### 2.2. Model calibration, implementation, and analysis

Calibration of the model began by finding previously published experimental data (4,30). We fit our mathematical model strictly to time course data using CaliPro, modified for this system (31). Fitted parameters were compared to previously published measurements of similar parameters, obtained through a literature search, in other systems to determine reasonable parameter ranges.

The model was implemented in SimBiology within MATLAB 2022a. Differential equations were solved using ode23s. The simulation was initially run for a long time (24 h) to find a basal steady state, following which induction of 17β-estradiol was simulated and the simulation continued (until 48 h) to obtain the new steady state. For data fitting, we used peak induction of the experimental data as the system steady state value.

The time course data and 24-h dimethylsulfoxide data from Brennan et. al. (30) were extracted using PlotDigitizer (32). For each data point, this yields a range between “high” and “low” values based on the error bars. Three data points – the basal steady state data point, and the timepoints at 24 h, 24+8 h (8 h post induction) and 24+24 (24 h post induction) – were fed into CaliPro. Model simulations were compared with the experimental data using a pass criterion. The initial pass criterion allows for ± 500% model output relative to the experimental data; after two iterations, the pass criterion is reduced to ± 200%; after four total iterations it is ± 50%, after 6 iterations it is ± 35%, after 8 iterations it is ± 25%, and, after 10 iterations, it is ± 10%. Once the model reaches this final pass criterion, when the number of pass parameter sets reaches 75% of the sampled 500 parameter sets per iteration, the calibration is designated as “complete”. Alternative Density Subtraction was used to narrow parameter ranges within CaliPro (31). The final parameter sets were then stored. From the final passed parameter sets, one set was randomly chosen and used for further analysis. Comparison of the model results and separate dose-response experimental results was done in MATLAB, using the curve fitting toolbox.

### 2.3 Model sensitivity analysis

Individual parameters were varied by 2x, 3x, 1/2x, and 1/3x and compared to the calibrated model output to determine which parameters might affect key assay attributes of dynamic range and sensitivity (Figure S5). Additionally, the initial conditions of state variables within the model were varied to study their effect on model output (Figure S5).

Global sensitivity analysis (GSA) was performed using the Sobol method and Sobol sampling (33). One thousand samples were used; increasing sample size did not affect the results shown in Supplementary Table S2. All simulations and analyses were further analyzed using developed scripts. All scripts are available on request from the authors.

## 3. Results

### 3.1. Genetic circuit model

Estrogenic transcriptional activation circuits have a similar architecture, shown in Figure 1, which serves as the basis for the mathematical model developed here (4– 8,34). A genetic circuit inserted into the cell line contains an estrogen response element (ERE), a minimal promoter element, and a reporter gene, typically luciferase. These circuits are based on transcriptional activity of a reporter gene following estrogen binding and activation. Estrogen receptors can be either endogenously (4), or exogenously expressed (5,7).

**Figure 1.**
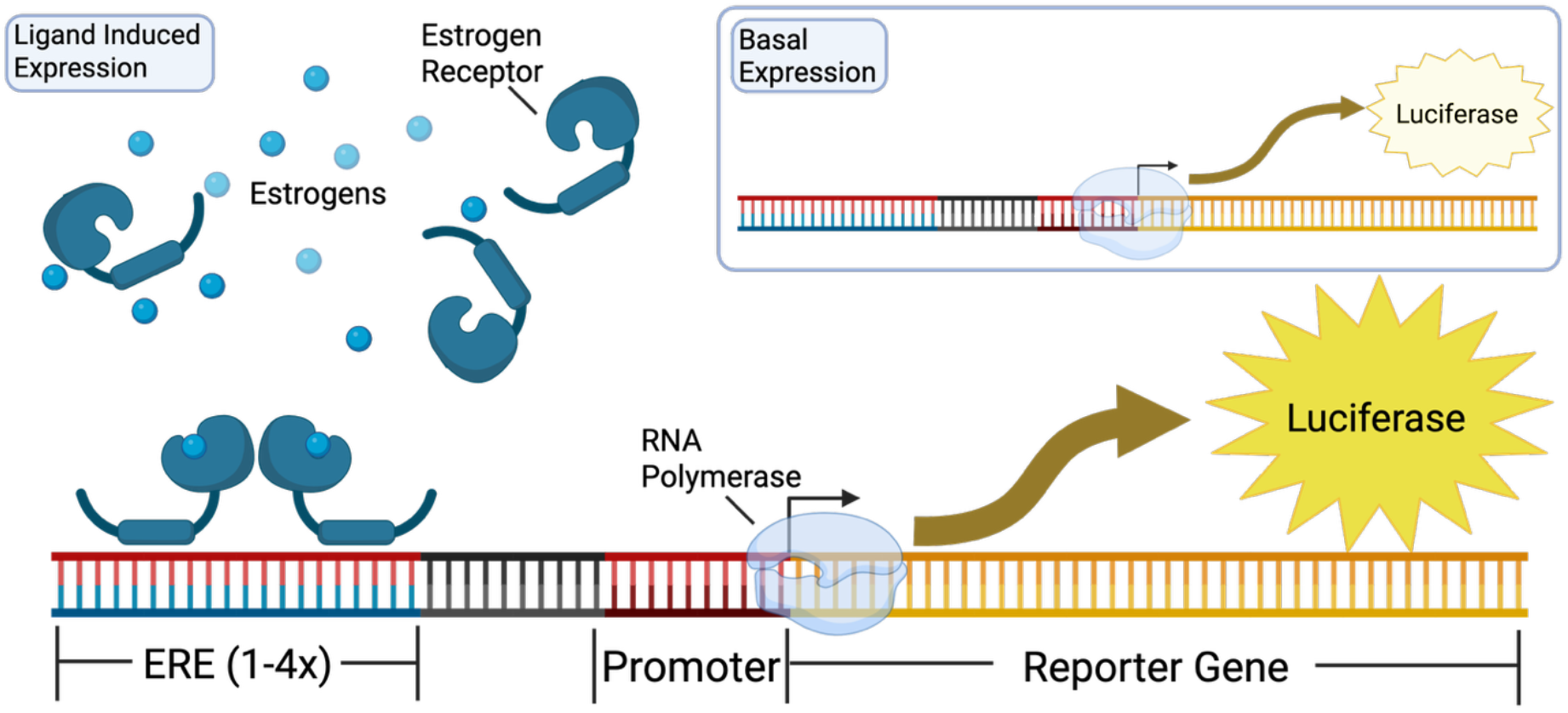
Schematic of the basic circuit common to reporter cell lines for estrogenic activity. A synthetic reporter cassette consisting of one or more estrogen response elements (EREs), a promoter region, and the luciferase reporter gene is inserted in a cell that expresses an estrogen receptor. An estrogenic chemical binds to the estrogen receptor, activating it. Activated estrogen receptors dimerizes and translocates to the nucleus where it binds to the EREs. RNA polymerase is recruited, and transcription of the luciferase reporter gene occurs. Luciferase mRNA is translated, the protein folds, and is expressed. In the absence of the ligand, a small level of basal expression produces a background signal.

The mechanisms of this circuit were modeled by a system of ODEs (Equation S1 and Figure S1) as following the law of mass action. An estrogenic chemical or ligand interacts with an ER, which then dimerizes, translocates to the nucleus, and binds to the ERE. Polymerase may be recruited with or without contribution from ligand-estrogen receptor complex. Polymerase activity leads to transcription of mRNA, then translation, and finally reporter protein (luciferase) synthesis. The units of each state variable of the model are molecules. Our ODE model consists of 11 differential equations containing 14 parameters.

### 3.2. Model calibration

Calibrating a model requires some basic knowledge of the molecular-level parameters. To initialize the model, we found plausible measured values from the literature. The kinetics of ER binding affinity, ER dynamics, and luciferase decay have been well studied and their parameters experimentally measured (35–37). Similarly, parameters related to mRNA and protein production in mammalian cells have been globally quantified and extensively studied (38). However, several parameters in the model are either difficult to accurately quantify or have not been studied. All parameters used for initialization are presented in Supplementary Table S1.

Initial parameters were sampled within a uniform distribution (Supplementary Table S1) and the system of equations was modeled. The first half of the simulation models basal expression. This part of the simulation assumed the output of luciferase reached a steady state by 24 h in the absence of estrogenic ligand and was calibrated to vehicle control data. Next, estrogen was added into the model system at a concentration of 0.1 nM and the modeled luciferase output at 8- and 24-h post-induction were then used for calibrating the model to experimental data. This strategy calibrates the model parameters related to basal expression and ligand-induced expression independently, specifically using time course data only.

Pass set criteria were used to evaluate the model outcome for any given set of parameters and to distinguish which parameter sets are used in the next round of model calibration (Figure S2). Finally, the distribution of parameters for the final model calibration, at which point >75% percent of the simulations were within 10% of both the first and second steady states, were stored for future analysis (Supplementary Table S1).

We compared fitted parameters to previously published measurements of similar parameters in other systems, obtained through a literature search, to determine reasonable parameter ranges (Table 1). We found that predicted parameters were reasonable and fell within measured ranges, with one exception: the half-life of luciferase, the reporter protein. Our parameter predictions suggested a much shorter luciferase half-life than the value reported (37), shown in Table 1.

**Table 1.** Literature values used for model calibration and the resulting calibrated model range.

| Parameter | Literature Value or Range (Units) | Model Value or Range (Units) |
| --- | --- | --- |
| ERE Number | 4 (–) (4) | 4 (–) |
| Estrogen Receptors | 260000 (–) (36) | 260000 (–) |
| Polymerase Molecules | 86400 (–) (39) | 86400 (–) |
| Estrogen Binding to Receptor (On) | $1.3 \pm 6 \times 10^6 \text{ (M}^{-1}\cdot\text{s}^{-1})$ (35) | $6.85 - 7.50 \times 10^6 \text{ (M}^{-1}\cdot\text{s}^{-1})$ |
| Estrogen Binding to Receptor (Off) | $1.2 \pm 2 \times 10^{-3} \text{ (s}^{-1})$ (35) | $1.01 - 1.04 \times 10^{-3} \text{ (s}^{-1})$ |
| mRNA Production | $0.1 - 100 \text{ (h}^{-1})$ (38) | $65.2 - 77.1 \text{ (h}^{-1})^1 / 17.3 - 17.4 \text{ (h}^{-1})^2$ |
| Luciferase Folding | 1-14 (min) (40) | 7.7 – 8.2 (min) |
| mRNA Decay (half-life) | 10 – 50 (min) (38) | 12.9 – 15.0 (min) |
| Luciferase (half-life) | 120 (min) (37) | 0.16-0.22 (min) |

<sup>1</sup> Value represents the range of ligand-induced expression rates
<sup>2</sup> Value represents the range of basal expression rates

The sets of parameters generated from calibration, stored as each individual sampled parameter set, were used to randomly select a single parameter set to use for further analysis. It should be noted that the two points occurring at 26 and 28 h (2 and 4 h after induction with 17β-estradiol) were not used in the model calibration in Figure 2(a) since we were interested in capturing the steady-state responses as these are most relevant for the assay.

**Figure 2.**
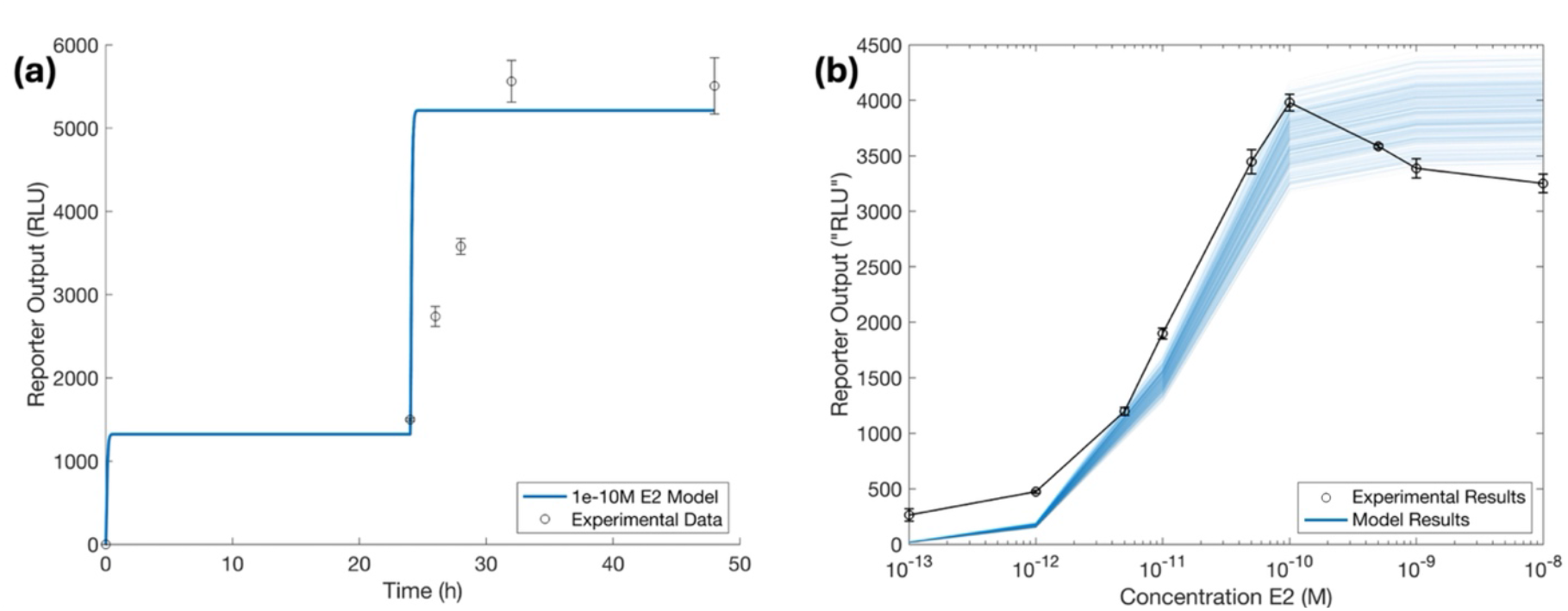
Model calibration and evaluation. **(a)** Model calibration, shown here using a randomly sampled passed parameter set, was evaluated against time course experimental data. **(b)** Steady-state values of all model outputs used for calibration were background-subtracted and plotted against experimental results developed in a separate dose-response experiment. All experimental data are from published literature.

### 3.3. Dose-response behavior recapitulated from time course calibration data

No further fitting of the model was conducted beyond what was done using the time course data. A random parameter set from the passed parameter set space was chosen. This parameter set was compared to dose-response experimental data by varying the concentration of ligand introduced into the simulation at 24 h (Figure 2 and Figure S3). The molecules of luciferase were then converted to RLU to draw comparisons between modeling and experimental data (41). Model-predicted output of luciferase was then quantitatively compared against experimental values and the attributes of each dose response curve (Figure S4 and Table 2).

**Table 2.** Table comparing model results of a single random parameter set, shown in Figure 2(b).

| Response Attribute | Model output | Literature value (30) |
| --- | --- | --- |
| Max | 5546 | $5236 \pm 71$ |
| Min | 1492 | $1259 \pm 60$ |
| Fold Induction | 3.7 | $4.2^1$ |
| EC50 (M) | $1.5 \pm 0.2 \times 10^{-11}$ | $1.3 \pm 1.0 \times 10^{-11}$ |
| log(EC10) – log(EC90) | -11.55 to -10.11 | -11.53 to -10.39<sup>1</sup> |

<sup>1</sup> Values computed from data presented in (30).

Predictions using the fitted parameters graphically matched well with the experimental data shown in Figure 2b (30). The EC50 of the model, 1.5 ± 0.2 χ 10^-11^ M, is consistent with the EC50 of the experimental data, 1.3 ± 1.0 χ 10^-11^ M. Similarly, the extent of induction of the circuit in response to 0.1 nM 17β-estradiol indicates that the model captures the dynamic range of the circuit. The model and the literature data corresponded well regarding the EC10 and EC90, further showing that the response predicted by the model independently agrees with experimental data.

### 3.4. Parameters affecting dynamic range

From the dose response curves generated by systematically varying each parameter (Figure S5), six parameters were found to have the most effect on the dynamic range of the model output: ERE and promoter numbers, rate of transcription, rate of translation, rate of mRNA decay, and rate of luciferase decay. Apart from rate of transcription, each parameter was found to linearly affect the output of the model, though the magnitude and directionality varied.

The number of reporter cassettes is assumed to be 4 in the base model (7). As this is increased to 8 and 12, the protein output correspondingly increased two- and three-fold. Similarly, as the number of EREs and promoters, considered a single variable in the model, decreased to one-half and one-third, the protein output decreased by one-half and one-third.

The rate of transcription had the largest effect on the reporter output, with a two-fold and three-fold higher value increasing protein output by 2.3-fold and 3.6-fold, respectively (Figure S5). The rate of transcription, when decreased, followed the same trend and led to decreased protein output by 2.3-fold and 3.6-fold. As was the case for EREs and promoters, a higher rate of translation increased protein output linearly. The rate of mRNA and luciferase decay, when increased, decreased protein output with the same fold difference. Similarly, when decreased, the decay rate of mRNA and luciferase increased protein output proportionately.

### 3.5 Parameters affecting sensitivity

Several parameters within the model had an observable impact on the sensitivity of the genetic circuit, shifting the dose response curve left or right (Figure S5f-i, m, n).

Parameters of the model that increased sensitivity are the affinity of ER to ERE and the rate of recruitment of polymerase.

Increasing the affinity of ER to the EREs by increasing the on-rate or decreasing the off-rate shifted the dose response of the simple circuit left (more sensitive) as shown in Figure S5(f, g). Interestingly, increasing on-rate and decreasing off-rates both had nonlinear effects. If the on-rate was increased 2x, or the off-rate was decreased by 0.5x, there was a 25% decrease in the EC50. If, on the other hand, the on-rate was decreased by 0.5x, or the off-rate was increased 2x, we observed a 30% increase in the EC50. This nonlinear trend extended to any change in the on-or off-rate parameter for ER binding to ERE (Figure S5(f, g)).

Recruitment of polymerase to the promoter occurred at different rates depending on whether it was promoted by ligand-binding to the ER or was ligand-independent. The effect of varying the rates of ligand-independent recruitment of polymerase displayed the same behavior as varying ER to ERE affinity (Figure S5h-i, m-n). The forward parameter displayed a non-linear relationship with the output, and the inverse parameter mirrored the exact same response in the opposite direction. Ligand-dependent recruitment of polymerase was unique (Table S3), as for this parameter luciferase output increased linearly as the on-rate increased, but changing the off-rate had almost no effect. Increasing or decreasing the off-rate by two-fold changed the EC50 by less than 5%.

### 3.6. Parameters exhibiting output insensitivity

Several parameters did not lead to output changes when their values were changed by 2-fold or 3-fold (Figure S5). The model output did not change with ER number, polymerase number, ligand affinity to ER, or the folding rate of luciferase. The number of ERs and the number of polymerase molecules also had no significant effect on the model output of luciferase (Figure S5a, c). Furthermore, the binding strength of estrogen, or the ligand of interest, to the ER as reported by the on- and off-rates had no observable effect for small differences (Figure S5d, e). Finally, the folding rate of luciferase had no observable effect on the output of luciferase (Figure S5l).

### 3.7. Global sensitivity analysis of parameters relevant for reproducibility

To test if results of single-parameter changes reported in Section 3 were robust when multiple parameters were changed simultaneously, we used GSA with Sobol sampling and identified parameters to which the output of the signaling circuit is most sensitive (Figure 3). This sensitivity analysis assumed a range of 10% in each parameter. While 10% was arbitrarily chosen, sensitivity analyses at other ranges returned similar results.

**Figure 3.**
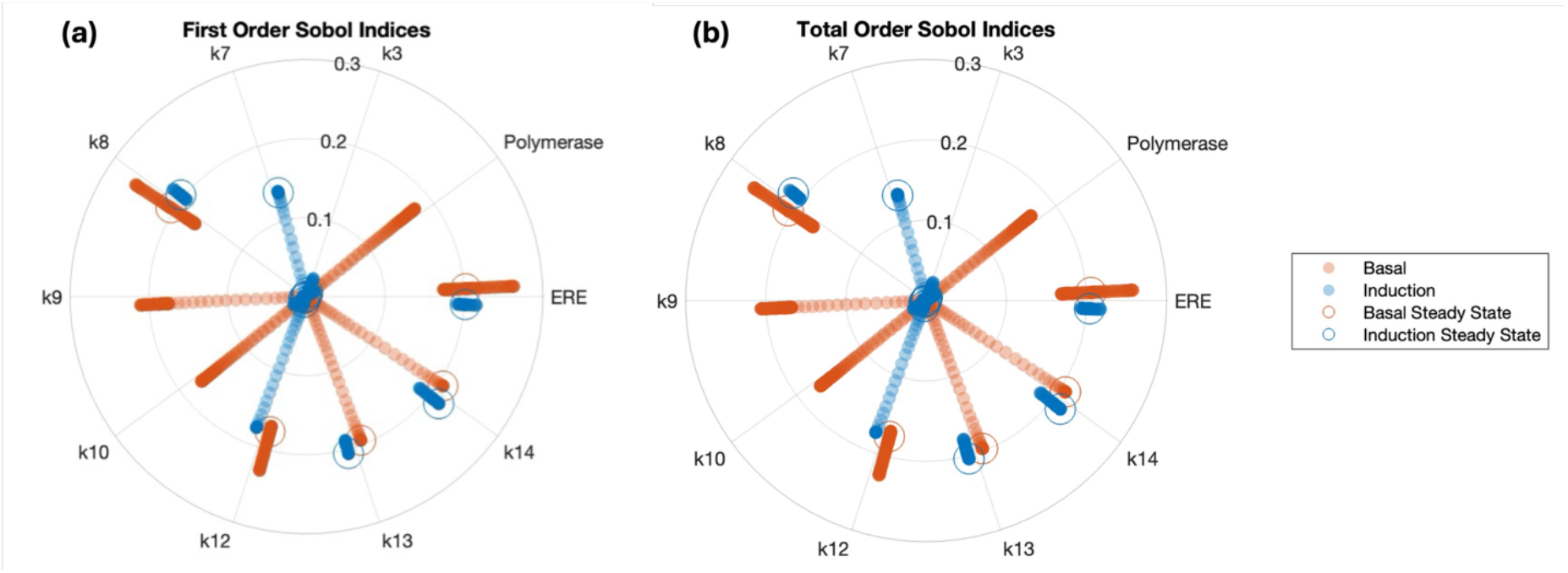
Global sensitivities of the model to parameter changes. (a) first order and (b) total order. The parameter range was 10% for the calibrated model evaluated with 0.1 nM 17β-estradiol (E2) added at 24 h. Orange points represent the sensitivity of the parameters prior to E2 addition, blue values represent the sensitivity of the parameters following E2 addition (induction), and circles represent their respective steady-state values.

Performing GSA by using a 10% range independently verified the results presented in Section 3.4 and 3.5 and shown in Figure S5. ERE and promoter number, k7 (rate of transcription), k8 (rate of translation), k13 (rate of mRNA decay) and k14 (rate of luciferase decay) were identified as having a significant effect on dynamic range. Steady-state Sobol index values are circled in Figure 3. When compared to previous results in Sections 3.4 and 3.5 regarding dynamic range and sensitivity, the use of GSA successfully identified parameters that most influence the model and may be targets for tuning the genetic circuit. GSA did not capture some parameter-reporter relationships, however. Parameter k3 and polymerase were identified as influencing sensitivity, but the steady-state values of their Sobol indices were relatively small (Figure 3).

Comparisons between first-order and total-order sensitivity indices reveal identical results (Figure 3a and 3b). The lack of a difference indicates that model parameters are not correlated with each other. Differences between the first-order and total sensitivity indices typically suggests that interactions between model parameters exist (41).

## 4. Discussion

We apply techniques from systems and synthetic biology to a subset of reporter gene assays used for toxicology, specifically assays used for detecting estrogenic chemicals. We propose parameters to tune the dynamic range and sensitivity of an existing reporter gene circuit by varying the model parameters and simulating the resulting dose-response curve. In addition, we used global sensitivity analysis to identify the parameters that most influence the gene circuit output based on this model. We predict that tuning the parameters highlighted from global sensitivity analysis would minimize variability of the reporter gene assay output.

CaliPro was selected for calibrating this estrogen receptor sensing circuit model due to its ability to function with systems that have experimentally unvalidated parameters. Unlike optimization approaches that minimize objective functions or Bayesian methods that require prior parameter estimates, CaliPro enabled exploration of a wide parameter space with minimal input requirements (31). Alternative methods like genetic algorithms and nonlinear regression failed to accurately capture time-course dynamics (data not shown). CaliPro was used to sample numerous global parameter sets and retained those meeting distance-based criteria from experimental data (31).

This approach allowed for selective parameter constraints and proved particularly valuable for developing this mechanistic model with incomplete experimental validation. The parameter sets are not true values – rather approximate the parameters specific to our model. The final model presented is not unique – refining or modifying a biological model is an iterative process – but this version serves as a starting point for experimentation and to demonstrate the sensitivity analysis reported here.

GSA allows for the identification of input-output relationships in models (33,42). Sensitivity analysis has been applied to provide mechanistic insight to applied models and is a powerful tool for model selection (44–46). GSA has also been used to identify potential targets for tuning a genetic circuit, as demonstrated with a genetic inverter circuit in *E. coli* designed to repress expression of a fluorescent protein (26). This demonstrates the capabilities of GSA for tuning genetic circuits and more generally represents the applicability of this type of model analysis on engineered biologic constructs. Here, sensitivity analysis indicates potential targets that have a substantial impact on the reporter protein output within the mechanistic model. Temporally dividing the sensitivity analysis allows for unique interpretation of parameters that affect the basal and ligand-induced expression separately, but the model does not assume correlation between these two cases. While GSA reports on sensitivity, additional validation by simulations is required to demonstrate how exactly parameter changes affect assay performance.

The modeled dose-response curve requires multiple simulations of the model at varying concentrations of ligand. It is not obvious that a dose-response curve would emerge from the model using varying ligand concentrations based solely on a calibration of the model to one ligand concentration with distinct time course data. This dose-response model that closely matches the experimental results suggests that the model captures the physical behavior of the gene circuit well. Manually adjusting parameters is a crude method to determine their effect; however, in this example, it is extremely useful. Local or global sensitivity analyses do not capture the effects of changing parameters on the dose-response curve, only on the time-course model. Also, compared to experimentally screening the effects, it is easier to computationally manipulate parameter values to predict their effects and validate those predictions with targeted experiments afterwards.

Analysis of the model revealed several parameters to target through design and experimentation for tuning the sensing circuit for estrogenic chemical screening. For example, several parameters could be targeted to tune dynamic range. The parameters that affected dynamic range the most were ERE and promoter number, rate of transcription, rate of translation, rate of mRNA decay, and rate of luciferase decay. Increasing the transcription binding factor site concentration has been shown to affect signal output in gene circuits (23). Tuning the rates of transcription and translation requires modification to the reporter protein gene, and there are limited options to do this without affecting the function of the reporter protein. The rates of transcription and translation are also dependent on concentrations of housekeeping proteins and the metabolic state, and can vary from cell to cell due to extrinsic noise (47,48). Modulating the stability of mRNA (or the mRNA decay rate) has been shown to affect steady-state protein levels (49) and is used in synthetic circuit design (50,51) but to our knowledge has not yet been used in assay design. Manipulating the decay rate of the reporter protein luciferase was shown to have a significant effect on dynamic range. Several methods exist for targeting reporter protein decay rate (52). However, since we do not explicitly model the luciferase light reaction, the decay rate of luciferase, k14, is a lumped parameter. Previous work in modeling and experimentation surrounding luciferase activity has led to insights into its complexity (37,53,54), and assumptions such as excess luciferin and ATP, along with co-factors, are required. Thus, our treatment of the reporter output may be an oversimplification, and further refinement of the model in this area may be necessary. It should be noted however that all parameters affecting dynamic range may also affect basal expression as an unintended consequence.

Similarly, the sensitivity of the modeled gene circuit appears to be tunable via parameters such as the binding affinity of ER and the recruitment rate of polymerase.

For an assay application, tuning the affinity of ER is a viable option but may lead to issues with non-specific binding. Recruitment of polymerase is a complex biological process and to our knowledge has not been targeted in current transcriptional sensor design. However, there are challenges in implementation since transcription factor binding elements designed too close to transcriptional start sites have been shown to downregulate gene expression (55).

These analyses together offer unique insights into the mechanism behind a simple genetic circuit. Tuning the most minimal sensing circuit will provide a foundation for other more complex circuit designs. Multi-parameter optimization along with circuit development including circuit repression may then lead to decreased basal expression while increasing dynamic range. This will ultimately provide a higher signal-to-noise ratio for the gene circuit. Coupling time-course data with mechanistic models and uncertainty analysis serves as a starting point for further circuit design. This saves time and resources by shifting experimental work to computational work, allowing for new hypotheses to be developed to tune sensitivity and dynamic range in genetic sensing circuits.

Given the enormity of the task of testing tens of thousands of chemicals for their effects on the estrogen signaling system, there is a need to develop new transcriptional activation-based cell assays that can supplement those already in use. To contend with issues of cellular heterogeneity that could affect reproducibility (15,16,56), forward engineering approaches that have become part of best practices in synthetic biology should be adopted. The strength of these methods is their ability to rationally identify parameters that contribute to performance and variability. This would help develop protocols to optimize performance while simultaneously minimize the effects of this variability, or alternatively, inspire the engineering of more sophisticated circuits to do the same (57,58). While this paper has focused on EDC detection, the workflow proposed here can be generalized to cell-based biosensors testing chemicals for activation of other signaling pathways.

## Supplementary Materials

## Author Contributions

AP, KFR and MG conceived research, MG and CH carried out research, MG wrote the first draft of the paper, AP, KFR and MG edited the paper, all authors read the final draft.

## Data Availability Statement

Matlab code used to analyze the data presented in the paper is available at https://github.com/prasadlabcsu/Estrogen-Receptor-Assay-Model

## Acknowledgment

We thank Connor Henderson and Daniel Hidalgo-Argote for their contributions by running simulations.

