## Supplementary Tables and Figures for "Quantitative modeling reveals sources of variability in transcriptional activation assays"

### Supplementary Information

#### Predicting Mechanistic Sources of Variability in Estrogen Receptor Transcriptional Activation Biosensor Assays

Mark Greenwood, Connor Henderson, Kenneth F. Reardon* and Ashok Prasad*
School of Biomedical and Chemical Engineering, Colorado State University

**Equation S1:**

d(Ligand)/dt = -(k1*Ligand*ER) + (k2*LigandBoundER) (1)

d(ER)/dt = -(k1*Ligand*ER) + (k2*LigandBoundER) (2)

d(LigandBoundER)/dt = (k1*Ligand*ER) - (k2*LigandBoundER) - (k3*LigandBoundER*ERE) + (k4*LigandERBoundERE) (3)

d(ERE)/dt = -(k3*LigandBoundER*ERE) + (k4*LigandERBoundERE) - (k10*ERE*Polymerase) + (k11*NoLigandPolBound) + (k12*NoLigandPolBound) (4)

d(LigandERBoundERE)/dt = (k3*LigandBoundER*ERE) - (k4*LigandERBoundERE) - (k5*LigandERBoundERE*Polymerase) + (k6*LigandPolBound) + (k7*LigandPolBound) (5)

d(Polymerase)/dt = -(k5*LigandERBoundERE*Polymerase) + (k6*LigandPolBound) + (k7*LigandPolBound) - (k10*ERE*Polymerase) + (k11*NoLigandPolBound) + (k12*NoLigandPolBound) (6)

d(LigandPolBound)/dt = (k5*LigandERBoundERE*Polymerase) - (k6*LigandPolBound) - (k7*LigandPolBound) (7)

d(mRNA)/dt = (k7*LigandPolBound) + (k12*NoLigandPolBound) - (k13*mRNA) (8)

d(UnfoldedProtein)/dt = (k8*mRNA) - (k9*UnfoldedProtein) (9)

d(Protein)/dt = (k9*UnfoldedProtein) - (k14*Protein) (10)

d(NoLigandPolBound)/dt = (k10*ERE*Polymerase) - (k11*NoLigandPolBound) - (k12*NoLigandPolBound) (11)

Equations used within MATLAB 2022a and within SimBiology in MATLAB2022a to model the estrogenic activity reporter circuit. The various species in the chemical reactions are defined in the network diagram shown in Figure S1.

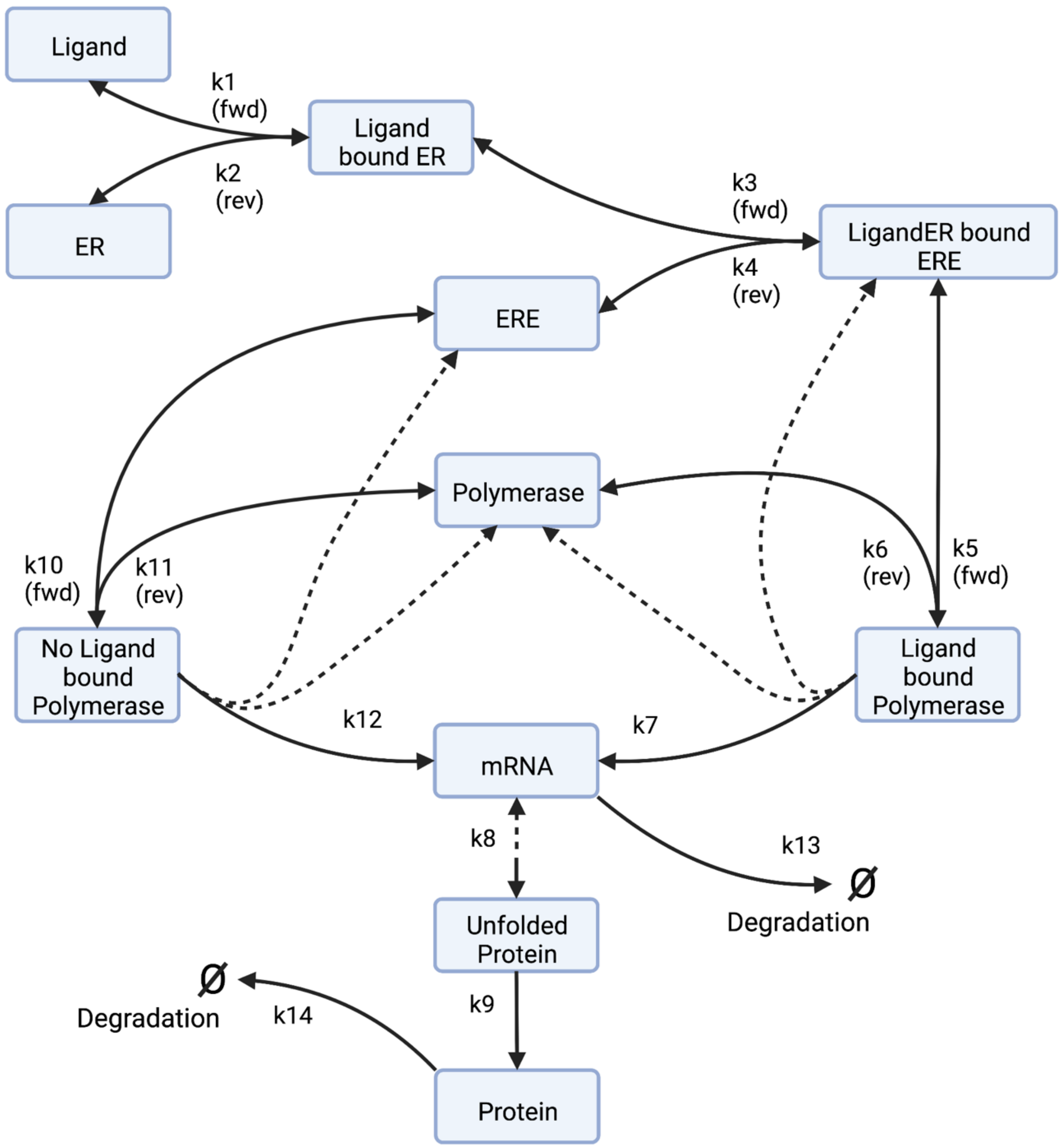

**Figure S1:** Model Diagram. Kinetic reactions are shown as yellow circles, state variables within the model are represented as blue ovals. State variables are further described as: Ligand – estrogenic ligand; ER – estrogen receptor; Ligand bound ER – Ligand-estrogen receptor complex; ERE – estrogen response element and promoter; LigandER bound ERE- Ligand-estrogen receptor-ERE complex; Polymerase – RNA polymerase; Ligand bound Polymerase – Polymerase interacting with Promoter with Ligand-ER complex bound to ERE; No Ligand bound Polymerase – Polymerase interacting with Promoter without Ligand bound ER. **—** Solid lines represent the direction of reaction. Single arrowed reactions imply that unidirectional flow is assumed while double arrowed reactions represent forward and reverse reactions. **- -** Dashed lines represent variables which are returned within a forward reaction, for example mRNA outputs unfolded protein, but is not degraded during the process. Created using Biorender.com

**Table S1**: Upper and Lower Bounds on all parameters used within model, shown before and after model calibration was conducted.

| Parameter Name | Lower Bound, initial | Upper Bound, initial | Lower Bound, final | Upper Bound, final | Units |
|---|---|---|---|---|---|
| k1 | 5.00E-06 | 6.00E-06 | 5.61E-06 | 5.64E-06 | $molecule^{-1} \cdot s^{-1}$ |
| k2 | 5.00E-04 | 1.80E-03 | 9.818E-04 | 1.001E-03 | $s^{-1}$ |
| k3 | 1.00E-05 | 1.00E-01 | 5.42E-02 | 5.47E-02 | $molecule^{-1} \cdot s^{-1}$ |
| k4 | 1.00E-04 | 1.00E+00 | 3.00E-01 | 3.26E-01 | $s^{-1}$ |
| k5 | 1.00E-08 | 1.00E-05 | 6.49E-06 | 6.78E-06 | $molecule^{-1} \cdot s^{-1}$ |
| k6 | 1.00E-08 | 1.00E-03 | 8.72E-04 | 8.96E-04 | $s^{-1}$ |
| k7 | 1.00E-03 | 1.00E-02 | 9.23E-03 | 9.24E-03 | $s^{-1}$ |
| k8 | 1.50E-01 | 5.00E-01 | 4.89E-01 | 4.90E-01 | $s^{-1}$ |
| k9 | 5.00E-04 | 3.00E-01 | 6.57E-02 | 6.94E-02 | $s^{-1}$ |
| k10 | 1.00E-08 | 1.00E-05 | 5.48E-06 | 6.10E-06 | $molecule^{-1} \cdot s^{-1}$ |
| k11 | 1.00E-04 | 1.00E-01 | 3.47E-02 | 3.49E-02 | $s^{-1}$ |
| k12 | 1.00E-05 | 1.00E-02 | 2.20E-03 | 3.22E-03 | $s^{-1}$ |
| k13 | 4.00E-03 | 1.27E-02 | 4.68E-03 | 4.83E-03 | $s^{-1}$ |
| K1e14 | 1.00E-05 | 1.00E-01 | 4.86E-03 | 5.58E-03 | $s^{-1}$ |

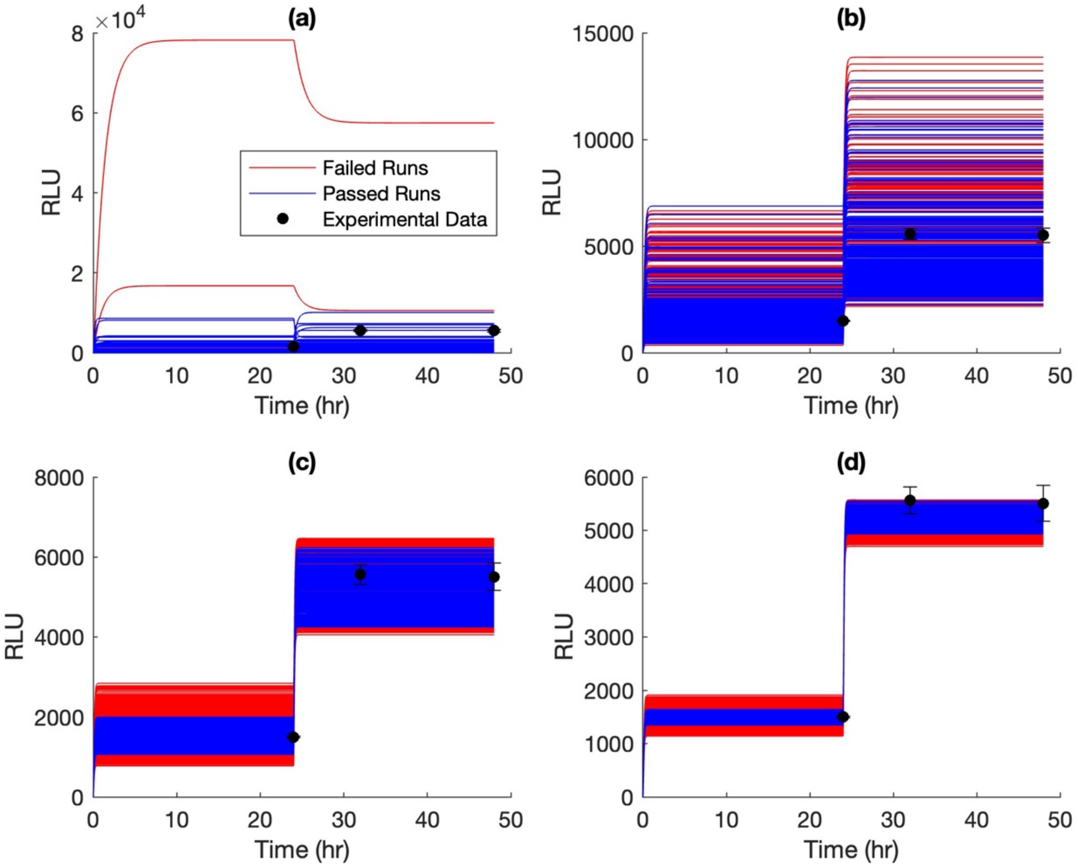

**Figure S2 (a-d):** All parameter sets identified by model calibration in (a) the first parameter selection, (b) the fifth parameter selection, (c) the tenth parameter selection, and (d) the final parameter selection. Passed parameters are shown in blue, failed parameters are shown in red, and experimental data from the supplementary of Brennan et. al., 2016 are shown in black dots with error [REF]. In (a), 4% of parameter sets passed with a flexible pass criteria within 2-fold of experimental data. In (b), 28% of parameter sets passed with a stricter pass criteria within 1.5-fold of experimental data. In (c), 37% of parameter sets passed, with the most strict pass criteria of 1.1-fold of experimental data, and finally in (d), greater than 75% of parameter sets passed. Code[1] in ‘passedvsfailedmodels.m’.

**Table S2:** Sobol indices from various sample sizes. The sample size used was 100, 1000, and 10000. Each sobol index for the timepoint at steady-state after ligand induction is shown.

| Sample Size | ER | ERE | Polymerase | k1 | k2 | k3 | k4 | k5 | k6 |
|---|---|---|---|---|---|---|---|---|---|
| 100 | 1.4E-06 | 2.2E-01 | -2.8E-04 | 2.3E-07 | -8.9E-07 | 2.2E-04 | 5.3E-03 | -1.2E-02 | -1.7E-06 |
| 1000 | 9.5E-08 | 2.0E-01 | 1.2E-04 | 2.2E-07 | 2.2E-08 | 9.0E-04 | 1.2E-04 | 9.1E-04 | 3.7E-06 |
| 10000 | 9.8E-09 | 2.0E-01 | 1.0E-04 | -3.1E-09 | 9.7E-09 | 5.0E-04 | 4.5E-04 | 9.1E-04 | 7.6E-06 |

| Sample Size | k7 | k8 | k9 | k10 | k11 | k12 | k13 | k14 |
|---|---|---|---|---|---|---|---|---|
| 100 | 2.0E-01 | 2.2E-01 | 6.9E-08 | 5.1E-05 | -4.2E-04 | -8.9E-04 | 2.4E-01 | 2.3E-01 |
| 1000 | 1.8E-01 | 2.0E-01 | -3.1E-09 | 3.9E-04 | 2.9E-06 | 1.3E-04 | 2.0E-01 | 2.1E-01 |
| 10000 | 1.7E-01 | 2.0E-01 | -3.2E-09 | 3.6E-04 | 3.0E-04 | 1.1E-04 | 2.1E-01 | 2.1E-01 |

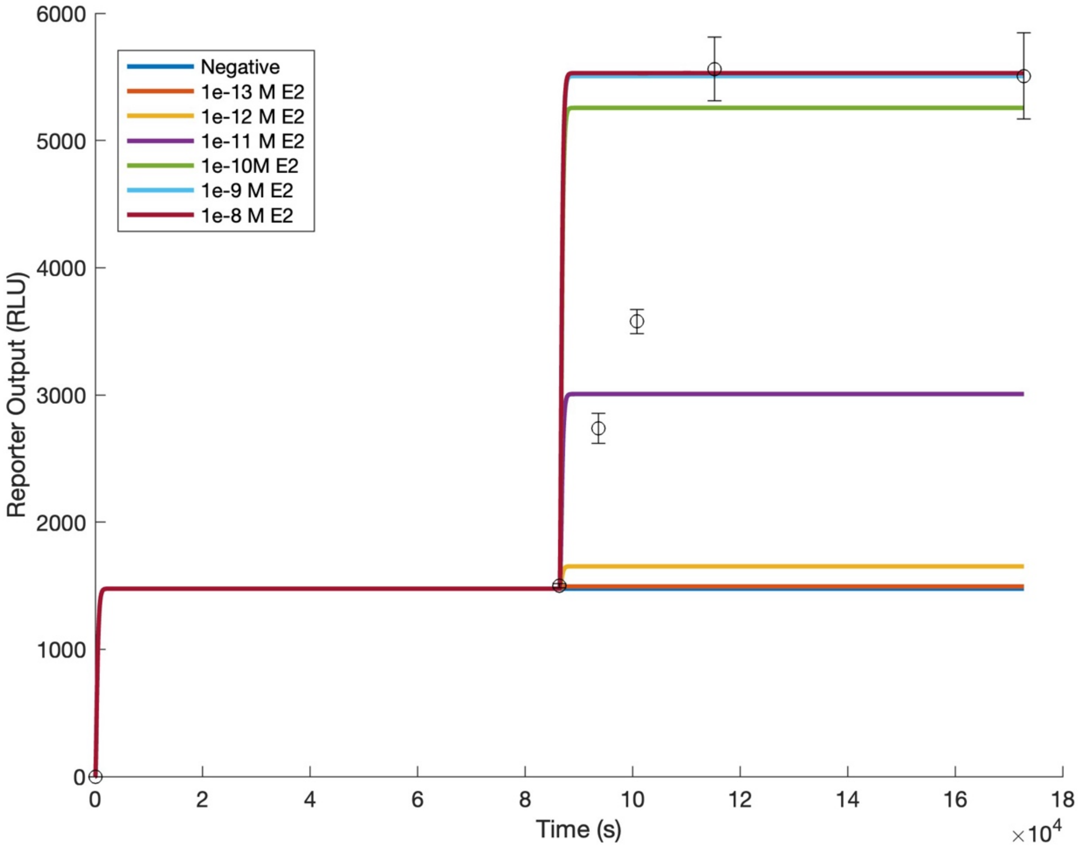

**Figure S3:** Time course data used to generate dose response curves. The steady state values at 172800 seconds, or 48 hours, corresponds to 24 hours post "induction" of a test chemical. Values were subtracted from the initial steady state at 86400 seconds, or 24 hours. Data points are shown in circles. Code[1] in 'DoseResponsePlot.m'.

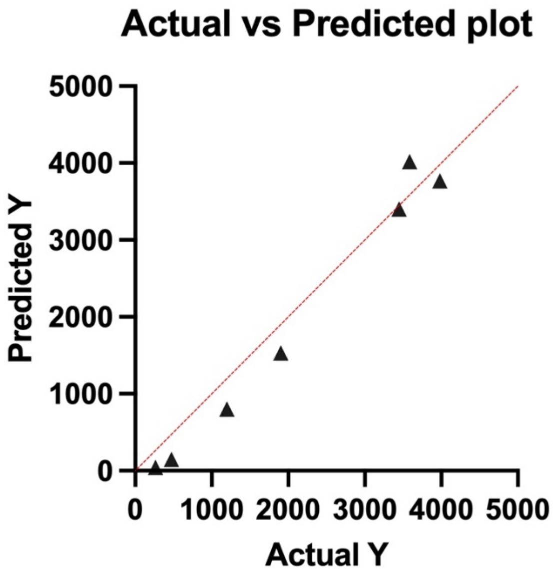

**Figure S4:** Plot of predicted modeled data (y-axis) versus actual experimental data (x-axis).

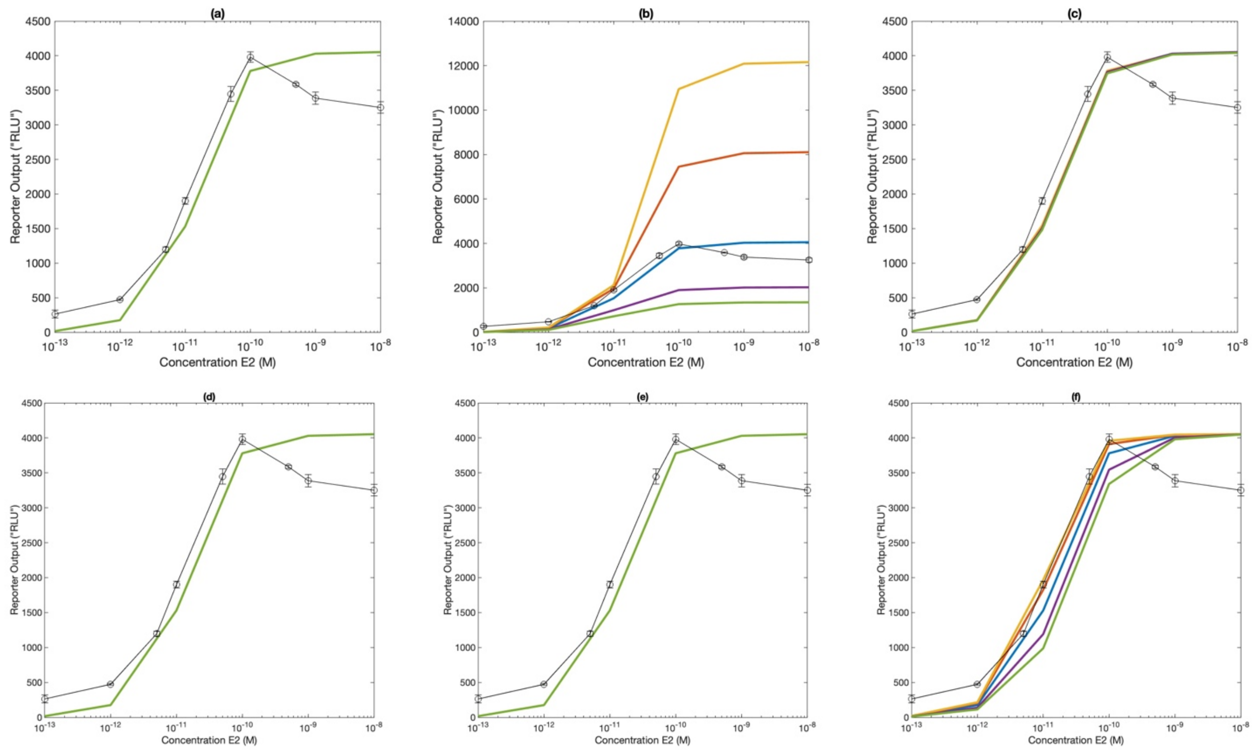

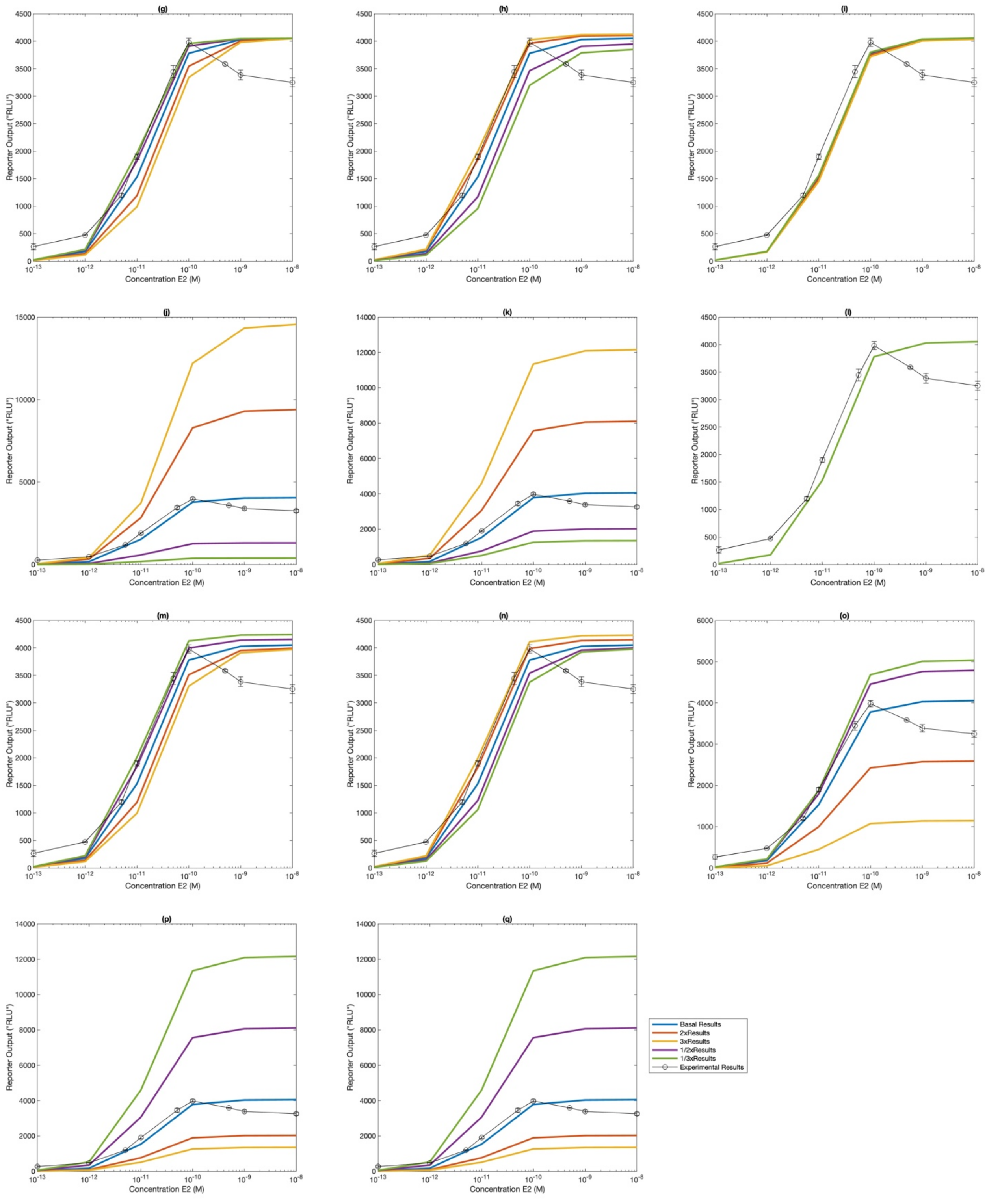

**Figure S5 (a-q):** Dose response plots, varying each respective parameter by 2-fold, 3-fold, 1/2-fold and 1/3-fold. Blue lines represent the original modeling results and black circles represent experimental data. Figure S5(a-c) shows results varying state variables "ER", "ERE", and "Polymerase". Figure S5(d-q) shows results varying all parameters, "k1" through "k14". So (d) k1, (e) k2, (f) k3, (g) k4, (h) k5, (i) k6, (j) k7, (k) k8, (l) k9, (m) k10, (n) k11, (o) k12, (p) k13,

(q) k14. Figures were made using code[1] in 'varymodelparam.m' and 'plotdoseresponsevarymodelparam.m'.

**Table S3**: Sigmoidal fit results from results shown in Figure S5(f, h, i, and m) investigating parameters which affect sensitivity of the output response. The factor shown indicates the amount to which only the single parameter was modified A 4 parameter sigmoidal equation using log[X] concentration values was used, with the R-squared of the fit shown. Key parameters, such as the normalized maximum and minimum values, hill slope of the plot, and IC 50 are shown. Results from parameters k4 and k11, shown in Figure S6(g and n), are not shown for brevity, but exhibited exactly the same response to the inverse multiplicative factor as k4 and k10, respectively.

| Parameter | **base** | **k3** | **k3** | **k5** | **k5** | **k6** | **k6** | **k10** | **k10** |
|---|---|---|---|---|---|---|---|---|---|
| Factor | - | 2x | 0.5x | 2x | 0.5x | 2x | 0.5x | 2x | 0.5x |
| Max | 4054 | 4063 | 4047 | 4114 | 3947 | 4044 | 4058 | 3992 | 4162 |
| Min | 39.43 | 50.69 | 23.94 | 51.23 | 23.74 | 37.62 | 40.44 | 24.42 | 51.19 |
| HillSlope | 1.341 | 1.435 | 1.227 | 1.434 | 1.230 | 1.328 | 1.349 | 1.233 | 1.428 |
| IC50 $*10^{-11}$ | 1.47 | 1.17 | 2.06 | 1.17 | 2.04 | 1.52 | 1.45 | 2.02 | 1.19 |
| $R^2$ | 0.9999 | 0.9998 | 1.000 | 0.9998 | 1.000 | 0.9999 | 0.9999 | 1.000 | 0.9998 |

#### REFERENCES

1. All code referenced here can be found at https://github.com/prasadlabcsu/Estrogen-Receptor-Assay-Model
